# Inactivation of SARS-CoV-2 in chlorinated swimming pool water

**DOI:** 10.1101/2021.04.19.440446

**Authors:** Jonathan C Brown, Maya Moshe, Alex Blackwell, Wendy S Barclay

## Abstract

SARS-CoV-2 transmission remains a global problem which exerts a significant direct cost to public health. Additionally, other aspects of physical and mental health can be affected by limited access to social and exercise venues as a result of lockdowns in the community or personal reluctance due to safety concerns. Swimming pools have reopened in the UK as of April 12^th^, but the effect of swimming pool water on inactivation of SARS-CoV-2 has not yet been directly demonstrated. Here we demonstrate that water which adheres to UK swimming pool guidelines is sufficient to reduce SARS-CoV-2 infectious titre by at least 3 orders of magnitude.

## Introduction

SARS-CoV-2, the causative agent of the COVID-19 pandemic, continues to transmit globally and makes quantifying the risks involved in different settings of great importance as societies attempt to return to normal. The potential for waterborne transmission of SARS-CoV-2 in the context of public swimming pools has not yet been investigated. Outbreaks of respiratory viruses such as adenoviruses, and enteric viruses such as enteroviruses, Hepatitis A and noroviruses which can transmit by the faecal-oral route are sometimes linked to swimming pools but often owe to improper maintenance of chlorine levels (Bonadonna & La Rosa, 2019; WHO, 2000). In the UK, swimming pools are treated with sodium hypochlorite to maintain a free chlorine level of 1.5-3 mg/l (ppm). The pH is also adjusted to between 7.0 and 7.6 as the availability of active free chlorine decreases with increasing pH (PWTAG, 2020). Here, by treating SARS-CoV-2 with swimming pool water which conforms to UK guidelines we demonstrate at least a 3-log_10_ reduction in infectious titre.

## Results

### Generating SARS-CoV-2 virus stocks suitable for inactivation testing

Virus stocks of SARS-CoV-2 for use in infectivity assays are generally generated by infection of a permissive cell line such as Vero and harvesting of virus in highly buffered cell culture medium. However, we observed in preliminary experiments that even a small amount of buffered medium was able to quench the chlorine activity of water samples whereas with an unbuffered saline solution the quenching was largely mitigated (not shown). The buffering capacity of the virus stock itself in cell culture medium would make it difficult to observe inactivation at the desired free chlorine and pH levels during testing. By infecting Caco-2 and Vero cells with a SARS-CoV-2 B.1 lineage virus at a multiplicity of 0.01pfu/cell, extensively washing off and replacing the growth medium with saline solution 24 hours before harvest at 3 days post-infection, we were able to generate stocks of infectious virus with reduced buffering capacity. To further minimise the effect of the non-viral constituents of the stock, such as cellular components which would exert a chlorine demand on the water samples tested, a 1:100 dilution of virus in normal saline was used in all inactivation tests.

### Inactivation of SARS-CoV-2 by chlorinated water

Water was collected from swimming pools in volumes of up to 1 litre and transported to the laboratory on the same day. The water was tested for free chlorine and pH levels upon arrival at the laboratory and adjusted to a range of values. A 1:100 dilution of SARS-CoV-2 virus stock generated in Caco-2 cells was then added to duplicate water samples in a total volume of 1 ml, incubated for 30 seconds at RT before quenching the chlorine with a one-tenth volume of 10X cell culture medium. Residual virus infectivity in the samples was then titrated on Vero cells by TCID_50_ assay. In each experiment the same virus stock was incubated for 30 seconds in PBS as a control.

Firstly, the effect of a range of increasing free chlorine levels in water starting at the minimum 1.5 ppm recommended in UK swimming pools were tested (Figure 1a). A low pH of approximately pH7 was used to give the best chance of observing virus inactivation as availability of free chlorine is maximized at lower pH. Under these conditions no detectable virus infectivity remained demonstrating at least a 3-log_10_ reduction in infectious titre compared to the PBS control where approximately 1×10^4^ TCID_50_/ml of virus was measured (Figure 1a). We next measured residual SARS-CoV-2 infectivity in conditions with higher pH while keeping the free chlorine level at approximately 1.5 ppm. Inactivation was observed to undetectable levels in all conditions except at the elevated pH of 7.62 at which low levels of virus infectivity were still observed at the threshold of detection of the assay. This inactivation equated to 3-log_10_ decreased infectivity compared to the control (Figure 1b). To demonstrate the interaction between the variables of pH and free chlorine in causing inactivation of SARS-CoV-2 infectivity, swimming pool water samples either at (1.42-1.72 ppm) or below (0.52-0.61 ppm) the UK recommended free chlorine levels were modified to pHs of approximately 7, 7.5 and 8. This resulted in only partial inactivation of the virus infectivity and revealed the importance of both chlorine levels and pH to achieve inactivation. (Figure 1c). Finally, we generated a further stock of the SARS-CoV-2 lineage B.1 virus in unbuffered saline in Vero cells and tested it against water at 3 pH levels at chlorine levels of 1.56 – 1.68 ppm. The new stock had a lower titre resulting in a yield of 1×10^3^ TCID_50_/ml from the PBS control condition. Nonetheless full inactivation equating to a greater than 2 log_10_ drop in infectivity was observed at pH7.00 and pH7.45, and even at pH7.80 the infectivity was decreased more than 50-fold (Figure 1d).

**Figure 1.**
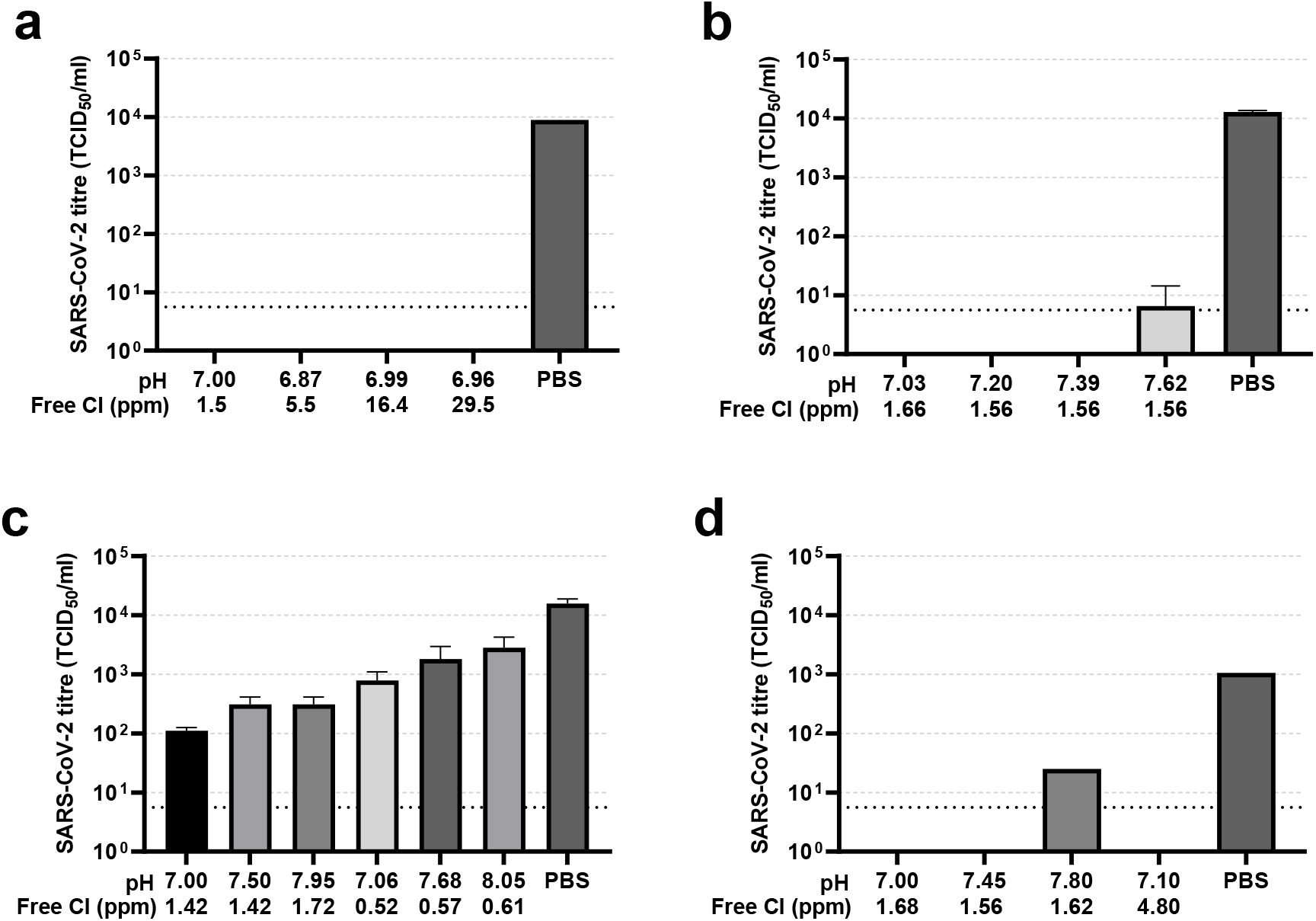
Exposure to chlorinated water inactivates SARS-CoV-2. Water samples taken from a swimming pool were modified in the laboratory to a range of pH and free chlorine values. A known amount of infectious SARS-CoV-2 was added to duplicate water samples in a volume of 1 ml, incubated for 30 seconds at RT and any remaining infectious virus then titrated by TCID_50_ on Vero cells. Residual virus titres are shown as the mean and SD of duplicate TCID_50_/ml values. Successive experiments were performed with varying free chlorine levels (a), varying pH (b), a range of both pH and free chlorine levels (c), and an independent preparation of virus at a range of pH and chlorine levels (d). A PBS control was included in each experiment to validate the infectivity of the virus input. Lower pH and higher free chlorine levels resulted in greater inactivation of SARS-CoV-2. A pH of no more than 7.4 and free chlorine above 1.5 parts per million (ppm) resulted in at least a 3-log_10_ reduction in viral titre.

## Discussion

Swimming pools have reopened in the UK as of April 12^th^ 2021 and therefore present locations of possible COVID-19 transmission. The likelihood of transmission events occurring in shared areas such as changing rooms can be minimised with social distancing and hygiene measures around the pool but different variables affect any risk associated with time spent in the water. Chlorination of swimming pool water has been used for decades to mitigate any onwards transmission of pathogens between swimmers. However, since the causative agent of COVID-19, the betacoronavirus named SARS-CoV-2, only emerged in late 2019, inactivation of SARS-CoV-2 by chlorinated water has not yet been directly demonstrated. Since viruses cannot replicate outside of a host, a transmission event via swimming pool water would require that virus emitted directly from a bather reached another at a sufficient infectious dose. Firstly, emitted virus will be greatly diluted before this occurs, potentially below a minimal infectious dose. In addition, if chlorinated water is directly viricidal against SARS-CoV-2, the likelihood of infectious virus being transmitted in swimming pool water will be further lowered. Demonstrating this may be important in increasing public confidence in returning to pools. Here we demonstrate that inactivation of SARS-CoV-2 in chlorinated swimming pool water is dependent on free chlorine and pH levels with increased inactivation at higher free chlorine and lower pH. We show that 30 seconds contact time at RT with water of a pH of no more than 7.4 and free chlorine above 1.5 mg/l (ppm) resulted in at least a 3-log_10_ reduction in viral titre (Figure 1). These levels are within the recommendations for swimming pools in the UK of at least 1.5 ppm free chlorine, although pH guidelines allow a pH of 7.0-7.6 and we found here that some residual virus was detected after treatment with water above pH7.4 even when at least 1.5 ppm free chlorine was present.

A limitation of this study is that we did not test survival of SARS-CoV-2 contained within mucus or saliva mixed with swimming pool water. Further we were only able to test reduction of a virus stock with infectivity around 10^4^ TCID_50_/ml due to the limited replication of SARS-CoV-2 in the laboratory and the need use a minimal volume of virus material during testing. Nonetheless, the viral challenge we presented equates to approximately 10^8^ genomes, (with a Ct value of 23) which is in excess of the amount of virus typically detected in the upper respiratory tract of asymptomatic people, with an average Ct of 31.15 (Ra et al., 2021). The route by which any residual virus in swimming pool water might infect another swimmer is not clear. SARS-CoV-2 is transmitted in the air and also by direct inoculation. There is also a potential faecal-oral route of transmission for SARS-CoV-2 (Guo et al., 2021). Our findings on the susceptibility of SARS-CoV-2 to inactivation by swimming pool water underscore the importance for those who maintain swimming pools to adhere to UK guidelines for chlorination, and this should give confidence in the safety of bathers when in the water. Finally, we stress that swimmers should continue to adhere to locally recommended social distancing rules both in and out of the water.

## Methods

### Cells and viruses

African green monkey kidney (Vero) cells (Nuvonis Technologies) were maintained in OptiPRO SFM (Life Technologies) containing 2X GlutaMAX (Gibco). Human epithelial colorectal adenocarcinoma (Caco-2) cells were maintained in DMEM, 20% FCS, 1% NEAA, 1% P/S. SARS-CoV-2 linage B.1 isolate hCoV-19/England/IC19/2020 (EPI_ISL_475572) was diluted in cell growth medium and used to infect confluent cells at a multiplicity of 0.01 pfu/cell and incubated at 37°C, 5% CO_2_. Growth medium was removed 2 days post infection, the cell sheet washed twice with saline solution (ddH_2_O, 0.9% NaCl) and replaced with saline solution. After a further 24 hrs virus supernatant was harvested and clarified by centrifugation.

### Water samples

Swimming pool water samples were collected from pools in London, UK and tested upon arrival at the laboratory. Free chlorine and pH levels were tested using a MD 100 photometer (Lovibond) to the manufacturer’s instructions for tests in Figure 1a-c and a PoolTest 25 (Palintest) for the test in Figure 1d. Chlorine levels of the water samples were increased by addition of sodium hypochlorite and pH was increased by addition of sodium carbonate or decreased using sodium bisulphate before retesting. Inactivation experiments were performed within 30 minutes of water sample preparation to minimise decay of chlorine levels.

### Inactivation testing and titration of residual virus by TCID_50_ assay

Treatment of SARS-CoV-2 with water samples was carried out as described in the text. In short 10ul of virus stock was added to 990ul of water sample, incubated for 30 seconds at RT before addition of 110ul of 10X MEM. Titration of residual virus was performed by TCID_50_ assay on Vero cells using cytopathic effect as the readout for infectious virus. In short, a half-log_10_ dilution series of each sample was performed and 4 replicates of each dilution transferred to 96-well plates of Vero cells, incubated for 1 hr at 37°C, 5% CO_2_ and replaced with cell growth medium. After 4 days, cells were stained with crystal violet and scored for either an intact, stained cell sheet or the absence of cells due to virus-induced cytopathic effect. For each condition, the Spearman-Karber method was used to calculate the 50% tissue culture infectious dose (TCID_50_) of the residual virus.

## Acknowledgements

We would like to thank Water Babies, Swim England, SPATEX Foundation and RLSS UK for their help, support and funding for this work. Special thanks to Ian Ogilvie, Richard Lamburn, Ellen Meyer, Rachel Chalmers, John Lee, Hannah Smith and Jared Crockford for providing technical expertise.

## Notes

### Competing Interest Statement

The authors have declared no competing interest.

